# Dynamic fate map of hindbrain rhombomeres in zebrafish

**DOI:** 10.1101/2023.01.27.523420

**Authors:** Mageshi Kamaraj, Thierry Savy, Sophie Salomé Desnoulez, Nadine Peyrieras, Monique Frain

**Affiliations:** Laboratoire BioEmergences (USR3695/FRE2039), CNRS, Université Paris Saclay, 91198 Gif-sur-Yvette, France; Laboratoire Matière et Systèmes Complexes (MSC, UMR7057), CNRS, Université de Paris Cité, 75013 Paris, France

## Abstract

Understanding of the morphogenetic processes that underlie the patterning of the vertebrate hindbrain requires establishment of rhombomere (r) cell lineages. Using long term *in vivo* two-photon imaging of zebrafish transgenic lines and automated image processing tools, we provide a method to reconstruct the r2-r6 lineage trees from the onset of gastrulation through early neurulation. We provide a dynamic fate map of hindbrain patterning at single-cell resolution. We show that rhombomere progenitor domains are aligned along the anteroposterior (AP) and dorsoventral (DV) axes as early as the shield stage. Rhombomere progenitor domains show a segmental organization, parallel to the blastoderm margin that predicts the future AP order of hindbrain. The DV organization of rhombomeres is set by the segregation of the progenitors along the DV axis of the embryo. Progenitors located at the dorsal/medial part of the blastoderm form the ventral domain of the rhombomeres while the lateral progenitors constitute the dorsal part. Our study sheds light on the clonal origin of individual rhombomeres, spatial and temporal patterns of cell division and migration of rhombomere progenitors throughout the early steps of hindbrain morphogenesis.

## Introduction

During zebrafish gastrulation, dorsal neuroectoderm cells from both sides of the embryonic axis undergo convergence and extension (CE) towards the dorsal midline and form the neural plate. Cells of the neural plate engage in mediolateral cell intercalation and extend along the anteroposterior (AP) axis. This forms the neural keel structure and turns into a cylinder-like solid structure called neural rod. Its lumen opens at the apical midline of the rod and forms the neural tube (Schmitz, Papan and Campos-Ortega, 1993). The efficiency of convergence depends on Planar Cell Polarity signaling (Ciruna *et al*., 2006; Tawk *et al*., 2007; Quesada-Hernández *et al*., 2010) and requires coordination with the extracellular matrix and the underlying mesoderm (Araya *et al*., 2014; Araya, Carmona-Fontaine and Clarke, 2016). At late neurulation, the hindbrain region undergoes a transient segmentation process along the AP axis, which results in seven or eight repeated units called rhombomeres. They present as visible swellings around 18-20hpf in zebrafish (Moens and Prince, 2002). Each rhombomere is a cell lineage unit and is specified with a unique pattern of gene expression. This segmental pattern of the hindbrain is essential for the organization of eight of the twelve pairs of cranial nerves to determine their structure and function (Moens and Prince, 2002; Krumlauf and Wilkinson, 2021). It also guides the migration paths of cranial neural crest cells that emanate from the dorsal part of the hindbrain (Lumsden and Keynes, 1989).

A fate map of the zebrafish CNS, with various subdivisions of the brain at 6hpf and 10hpf, was obtained by the conventional method of sparse lineage labeling using fluorescent dyes (Woo and Fraser, 1995). This method allowed only a few surface-disposed cells to be followed over large time intervals. Tissue reorganization, cell cycle length and clonal strings during zebrafish neurulation were examined using the same method (Kimmel, Warga and Kane, 1994; Papan and Campos-Ortega, 1994; Woo and Fraser, 1995; Kozlowski *et al*., 1997). Cell movements and division orientation of surface cells of the epiblast were analyzed with time-lapse analysis (Concha and Adams, 1998). While these studies provided key insights into zebrafish CNS formation, there are currently no quantitative studies on the behavior of individual hindbrain progenitors during gastrulation through neurulation. Technical challenges have stood in the way of obtaining single-cell resolution. Further, the CNS fate map does not provide a precise fate map for the hindbrain with its individual rhombomeres. Establishing an accurate dynamic fate map of hindbrain rhombomeres should help decipher the cell rearrangements and cell behaviors that underlie its segmental organization.

Technical advances in microscopy, image processing and computational tools have enabled the reconstruction of cell lineages and cell behaviors of the intact zebrafish embryo for early stages of development (Keller *et al*., 2008; Olivier *et al*., 2010; Faure *et al*., 2016). The first study on reconstruction of zebrafish cell lineages focused on mesendoderm movements (Keller *et al*., 2008). A more recent study described the cell behaviors during germ layer formation, where global cell movements were examined (Shah *et al*., 2019). Other studies provided the organization of forebrain and spinal cord morphogenesis in zebrafish (England *et al*., 2006; Wan *et al*., 2019). Neural tube morphogenesis at the hindbrain has been described but only at late gastrula and early neurula stages (Hong and Brewster, 2006; Araya *et al*., 2019). Thus, we aim to analyze the behaviors of naïve ectoderm cells at single-cell resolution during their transition to neural plate and neural keel.

Our study aims to reconstruct cell lineages of hindbrain rhombomeres and assess cell behaviors such as proliferation and cell displacements. Using two-photon microscopy, we captured the formation of rhombomeres at single-cell resolution in a hindbrain reporter transgenic zebrafish, *krox20:eGFP-Hras*. By processing the imaging data through the BioEmergences workflow (Faure *et al*., 2016), we obtained an automated reconstruction of cell lineages of rhombomeres. After extensive manual corrections, we produced the first dynamic fate map of the hindbrain, from the onset of gastrulation through early neurulation, at single-cell resolution. We quantified the patterns of cell proliferation and cell movements of the rhombomere progenitors.

## Results

A predictable AP and DV organization in the presumptive neuroectoderm was demonstrated at the early gastrulation stage through sparse lineage tracing with a fluorescent dye (Woo and Fraser, 1995, 1998). However, the reported regional fate map of zebrafish CNS morphogenesis, established at two stages of gastrulation, fails to capture its dynamics and continuity. For these reasons and in order to understand how hindbrain patterning, cell fates and movements are coordinated during gastrulation, we performed a systematic study of hindbrain morphogenesis based on the reconstruction of cell lineages.

### AP segmental organization of rhombomeres’ progenitors at the onset of gastrulation

We generated zebrafish transgenic lines, Tg *krox20:eGFP-Hras* on the WT/Tü and *casper* backgrounds. *Casper* background allows the transgenic line to be devoid of pigmentation. *krox20:eGFP-Hras* drives cell membrane staining in rhombomeres 3 and 5 thanks to cA, an autoregulatory element of *krox20* expression (Chomette *et al*., 2006). This reporter line allows the identification of prospective rhombomere 3 and 5 domains upon their formation, with an hour delay compared to the initiation of endogenous *krox20* gene expression. Injection of *H2B-mCherry* mRNA at the one-cell stage into transgenic eggs provides ubiquitous nuclear staining, instrumental for automated cell tracking (Faure *et al*., 2016). We used two detection channels for emitted fluorescence, the eGFP (membrane) channel for identification of rhombomere cells and the mCherry channel (nuclei) for cell tracking. We established a 3D+time imaging protocol of zebrafish hindbrain formation from the onset of gastrulation through early neurulation using two-photon microscopy (Fig. 1A). We standardized the imaging parameters in order to capture complete hindbrain formation from its progenitor domains in the field of view with optimal spatial and temporal resolution required for cell tracking. The 4D image datasets were processed through the BioEmergences workflow using the Difference of Gaussian algorithm for image filtering and nucleus detection, and Expectation-Maximization algorithm for cell tracking, which provide automated cell tracking (Fig. 1B). Upon obtaining the reconstructed digital embryos, we superimposed the centers of the nuclei of the entire embryo on the membrane fluorescence channel, as eGFP membrane staining reports on the r3 and r5 domains. Nuclei centers of r3 and r5 domains in the digital embryo identified by membrane staining at 12hpf40 and 14hpf20, respectively, were marked with specific colors (Fig. 1C). Nuclei centers of the r4 domain (which lies between the r3 and r5 domains), were also marked. We extended our analysis to r2 and r6 whose nuclei selections are less accurate due to uncertainty about the location of the r1/r2 and r6/r7 boundaries. To define the spatiotemporal organization of rhombomere progenitor domains, we backtracked the r2-r6 selected cells till the onset of gastrulation (6hpf). Cell tracks were manually curated with the interactive visualization software, Mov-It (Faure *et al*., 2016). The number of selected cells in each rhombomere domain and the number of validated/manually corrected cells over the entire developmental period analyzed are given (Table 1). The automated reconstruction of cell lineages during early neurulation demanded more manual correction due to a lineage score of the workflow much lower than the 96% described for the first three hours of zebrafish embryonic development. The number of tracked cells from posterior rhombomeres, r5 and r6, was 50% less, as it was challenging to follow cells given the complexity of neural tissue at 14hpf. Nevertheless, we obtained the dynamic fate map of individual rhombomeres (Fig. 1C).

**Table 1.** The number of selected nuclei/cells in each rhombomere and the corresponding number of cells annotated till 6hpf.

| **Rhombomere** | **Selected cells** | **Annotated cells** |
| --- | --- | --- |
| r2 | 402 | 220 |
| r3 | 437 | 230 |
| r4 | 354 | 206 |
| r5 | 403 | 142 |
| r6 | 279 | 86 |

**Fig. 1.**
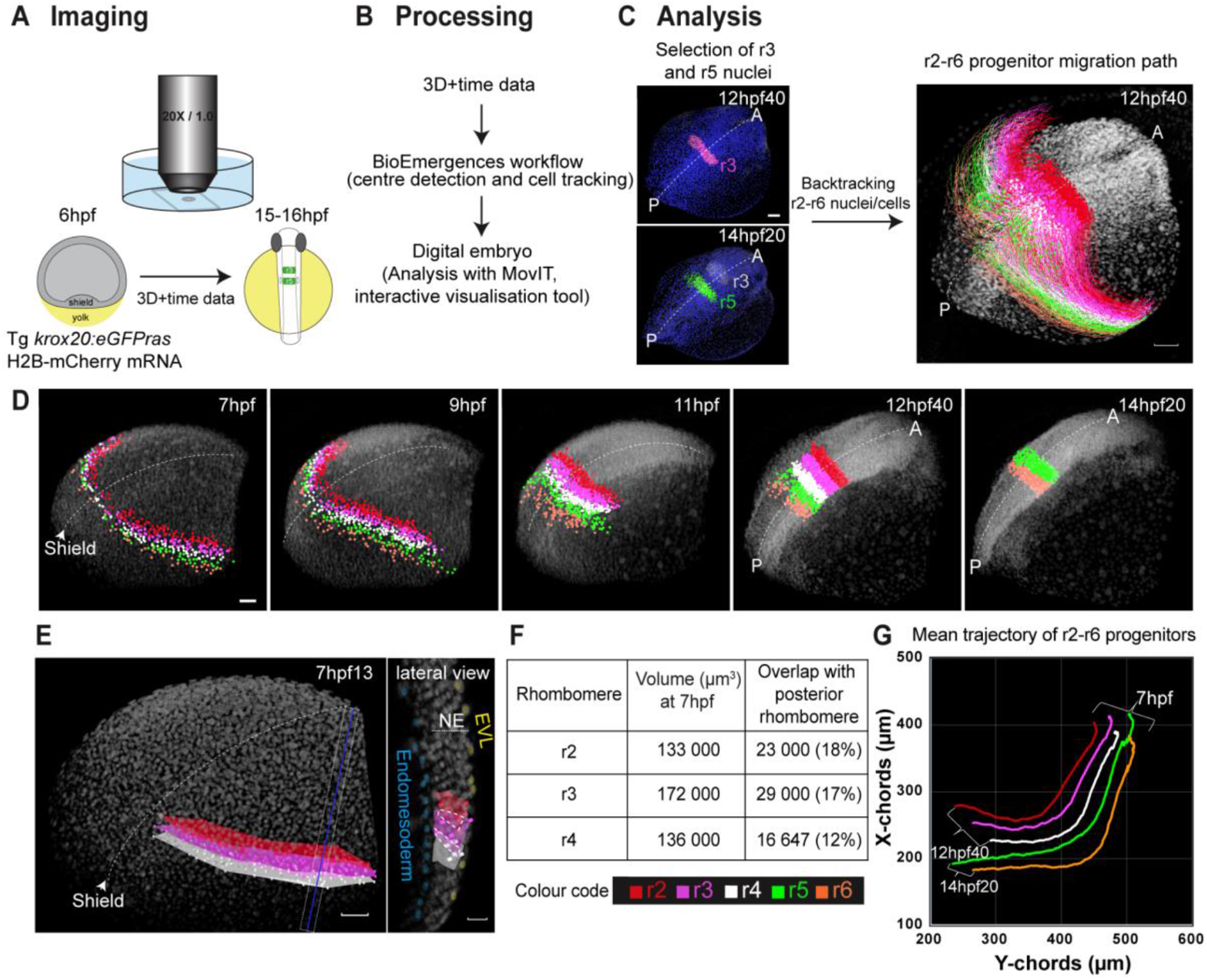
Segmental organization of rhombomeres’ progenitors is refined during gastrulation and early neurulation. A) Imaging scheme: 2-photon imaging of Tg krox20:eGFPras embryo injected with H2BmCherry mRNA from 6hpf to 15-16hpf. B) Processing of 3D+t imaging data through BioEmergences workflow for nuclei center detection and automated cell tracking. C-E) Analysis of the processed data, dorso-lateral view 3D rendering with nuclei staining raw data in gray, anterior top right (A), posterior bottom left (P), scale bar 50 µm except for E lateral view 20 µm, midline marked by a dotted white line. C) Left panel: detected nuclei centers in blue, selection of r3 (pink) and r5 (green) nuclei at 12hpf40 and 14hpf20 based on eGFP expression raw data in white. Right panel: tracks of r2-r6 nuclei from 6hpf to 12hpf40 displayed at 12hpf40. Color code: r2 red, r3 pink, r4 white, r5 green, r6 orange. D) r2-r6 progenitors’ nuclei position at 7hpf, 9hpf, 11hpf, 12hpf40. r2-r4 progenitors’ nuclei were not tracked beyond 12hpf40. At 14hpf20 display of r5-r6 nuclei only. E) Volume/surface estimation of r2-r4 progenitor domains on the right side of the embryo at 7hpf13. Dotted blue line indicates the plane of a transverse section 20 µm thick, 3D rendering (right panel). Dashed white boxes indicate r2/r3 and r3/r4 progenitor domains overlap. EVL: enveloping layer (detected nuclei yellow), NE: neuroectoderm, endomesoderm detected nuclei blue. F) Estimated volume of r2, r3 and r4 progenitors’ domain at 7hpf and overlap with the posterior neighboring rhombomere. G) Mean trajectory of r2-r6 progenitor population (right side of the embryo) plotted in the XY axis from 7hpf to 12hpf40 or 14hpf20 (r5-r6).

The position of r2-r6 progenitors was followed in the developing embryo from the onset of gastrulation through early neurulation (Fig. 1D; Movie 1; Fig. S1A). The hindbrain progenitors appear split into two domains on either side of the embryo midline. As shown for the precursors of the major brain subdivisions in (Woo and Fraser, 1995), we observed a segmental organization of r2-r6 progenitors along the animal pole-margin axis as early as the shield stage, which reflects the future AP order of the hindbrain. Using the following conventions, the center of the embryonic shield is defined as 0° longitude, the animal pole is defined as 90° latitude. We estimate that the r2-r6 progenitors are located between 10-100° longitude and 10-30° latitude which is similar to the published hindbrain precursor fate map (see Movie 1; (Woo and Fraser, 1995)). The progenitor population for each rhombomere occupies a distinct domain with a significant 12-18% overlap at the borders with adjacent progenitor populations at 7hpf (Fig. 1E). Some refinement occurs through formation of sharp boundaries at later stages. The volume of r2-r4 progenitor domains and their overlap with the next caudal progenitor domain were measured by calculating the volume of their alpha shape segmentation (Fig. 1F). We speculate that, at the onset of gastrulation, the clonal origin of rhombomeres is not yet established, thus allowing for greater mixing at the borders of adjacent rhombomere progenitor populations. Throughout the course of convergence and extension (CE) movements during gastrulation, we observed that rhombomere progenitors sweep posteriorly and then medially to form the neural keel/rod structures through collective cell displacements. This is illustrated by the mean trajectory (xy-axis) of r2-r6 progenitor populations from one half of the embryo plotted over time. We observe remarkably parallel and increasingly caudal paths, corresponding to the order of the respective rhombomeres in the neural tube (Fig. 1G, Fig. S1B). In short, the analysis of two Tg *krox20:eGFP-Hras* embryos from both sides (ID Datasets: 180516aF in Fig.1 and 190828aZ in Fig. S1) show that the predictable AP segmental organization of rhombomere progenitors is refined during gastrulation and early neurulation through a collective flow of cell movements with no or little mixing.

### Dorsal and ventral hindbrain progenitors are segregated

A significant dorsal and ventral order of the midbrain-hindbrain progenitors was previously reported for the 6 and 10hpf stages of zebrafish embryos (Woo and Fraser, 1995). Cells located medially in the neural plate internalize to form the ventral part of the neural tube. Cells located laterally constitute its dorsal part during the neural plate to neural keel/tube transition (Papan and Campos-Ortega, 1994; Araya *et al*., 2019). Here we explore the dynamics of DV segregation of entire progenitor populations of r2-r6 from the shield stage (6hpf) to the neural keel stage (13hpf). In the dorsal view of the reconstructed embryo, cell trajectories of r4 progenitors, taken as an example, showed two cell populations with distinct migration paths from 10hpf30 onwards (Fig. 2A). One population, labeled in white, moves posteriorly, then undergoes a flanking movement to join in the medial anterior flow and internalizes to form the ventral domain of the neural keel. The second population, labeled in yellow, converges directly to the midline to give rise to its dorsal domain. At 7hpf, the progenitors are segregated along the DV axis of the embryo. The ventral progenitors are located at the most dorsal part of the blastoderm, while the dorsal progenitors are located more laterally (Fig. 2A; Fig. S2A). Up to 10hpf30, these two populations move medially and anteriorly (converge) in the same direction toward the midline, then they follow separate paths (Movie 2). We next tracked the progenitor populations in the transverse plane (Fig. 2B, Movie 3). Between 10hpf30 and 11hpf20, the progenitor population close to the midline starts internalization in the neural keel as it forms. Between 11hpf20 and 13hpf, as the medial progenitor population moves deep into the neural keel, the lateral progenitor population continues to move toward the midline to end up on top. These cell trajectories recapitulate neural plate convergence at the midline as well as internalization to orchestrate the formation of the 3D neural keel structure (Fig. 2B; Movie 3). The mean trajectory (xy-axis) of the ventral and dorsal progenitors of rhombomere 4 (from one half of the embryo) summarizes their distinct paths in the embryo from shield stage to early neurulation (Fig. 2D and for r3 in second dataset in Fig. S2B). The same analysis was performed for the progenitors of others rhombomeres (r2, r3, r5 and r6), which show parallel paths for the dorsal and ventral subpopulations respectively, with a progressively more caudal localization that corresponds to the order of the rhombomeres along the AP axis of the embryo (Fig. S3A). Analysis of cell velocity showed that, between 8hpf40 and 10hpf40, the entire set of dorsal progenitors of r2-r6 transiently migrate faster toward the midline than the entire set of ventral progenitors (Fig. 2E). The same transient increase in cell velocity is observed for the dorsal progenitor population of rhombomere 4 when compared to the ventral population (Fig. S3B). This is in agreement with a previous report that showed an increase in speed of lateral cells, peaking between yolk plug closure (YPC, 9.5hpf) and the one-somite stage (10.3hpf)) (Sepich *et al*., 2000). Our analysis differs slightly from this study with respect to timing. We noticed that the dorsal progenitors of posterior rhombomeres migrate faster than their anterior counterparts (Fig. 2C). Overall, we provide a refined DV ordering of r2-r6 rhombomere progenitors in terms of migration paths and cell velocity.

**Fig. 2.**
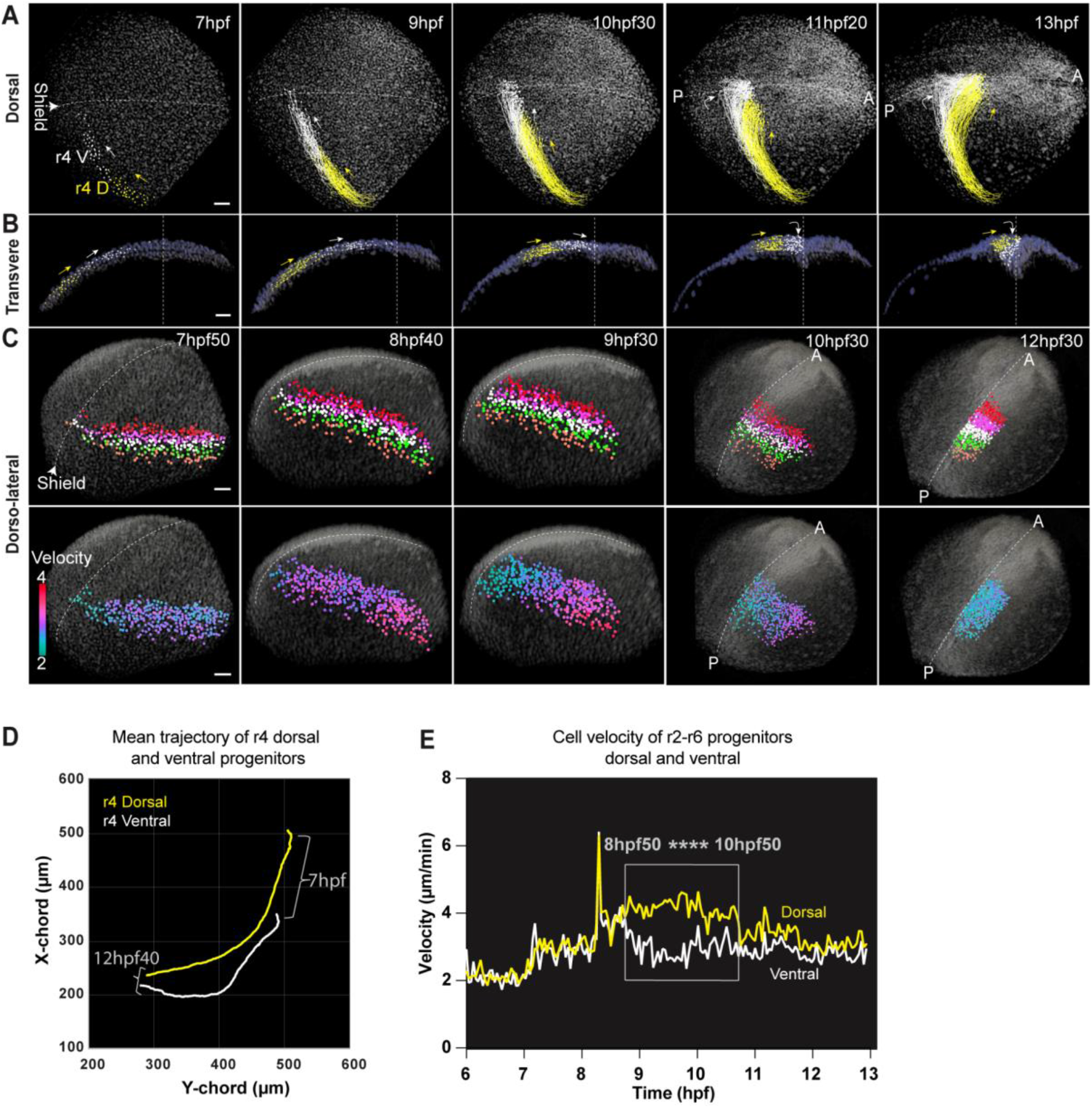
Dorsal and ventral hindbrain progenitors have distinct convergence/extension paths. A-C) 3D rendering of image data, nuclei raw data in gray, nuclei selected on the right side of the embryo with color codes for dorso-ventral domains in A-B, rhombomeres in C top panel as in Fig. 1, velocity in C bottom panel, time in hpf top left, midline indicated in white dotted, orientation indicated on the left, scale bar 50 µm. A) r4 progenitors’ position and trajectory from 7hpf to 13hpf, dorsal (D yellow) and ventral (V white), arrowhead points to the shield, arrows indicate the direction of progenitors’ migration. B) Data as in A except for tracks, here over 25 min, transverse section encompassing r4 domain (44 µm), ventral (white) and dorsal (yellow) progenitors, other detected nuclei in blue. C) Top panel as in Fig.1D. Bottom panel velocity map, color code from 2 to 4 µm/min. D) Mean trajectory of dorsal and ventral progenitors of r4 (right side of the embryo) plotted in the XY axis from 7hpf to 12hpf40. E) Relative velocity of dorsal (yellow) and ventral (white) progenitors of r2-r6 from 6hpf to 12hpf40. Mann Whitney test for the difference in mean velocity between 8hpf50 and 10hpf50 P<0.0001.

### The clonal origin of rhombomeres is set by mid-gastrulation

How is the hindbrain built as it undergoes transient segmentation during early embryogenesis? What is the contribution of cell proliferation in the formation of a segment versus the recruitment of cells from its neighborhood? To answer these questions, precise measurements are required. We therefore analyzed the clonal history of rhombomere progenitors by reconstructing the cell lineage trees of rhombomeres’ r2-r6 and exploring the fate of hindbrain progenitor cells. The lineage tree for each rhombomere from r2 to r6 was reconstructed by tracking the cells backward to their progenitor state and following their divisions (Fig. 3A and Fig. S4A). For each progenitor division we tracked the daughter cell forward and color-coded the daughter cell according to its localization in the segmented hindbrain. In the cell lineage trees of r2-r6 from the first dataset (ID:180516aF shown in Figs. 1 and 2), we examined at which stage clonal commitment of rhombomere progenitors occurs (r3 in Fig. 3A, r2-r6 in Fig. S4A). We marked the time point at which the progenitors began to give rise only to daughters of the same rhombomere fate. We found that the clonal origin of individual rhombomeres is achieved at around 8hpf15-8hpf40 (Fig. 3A). The commitment of neuroectoderm cells to neural fate and regional rhombomere identity occurs at 80% epiboly (8-8.5 h) as shown by microsurgical transplantations (Woo and Fraser, 1998). Over the 8 hours of embryonic period of development analyzed (6 to 14hpf), we found that 84-89% of progenitors divide into daughters of the same identity (Fig. S4B). But at early stages from 6 to 8hpf40 before clonal origin is achieved, we observed that 40-68% divisions contribute to adjacent rhombomeres (Fig. 3B). After the clonal origin is established, few subsequent divisions that occur predominantly at the rhombomere borders would contribute to the flanking rhombomeres due to some cell rearrangements i.e intercalation and internalization that take place during neural keel formation. These cells are known to undergo either cell sorting or a cell identity switch to make homogeneous segments with sharp boundaries (Xu and Wilkinson, 2013; Calzolari, Terriente and Pujades, 2014). During the 8 hours of embryonic development analyzed, cells undergo at least one and up to two cell division(s). Although progenitor cell divisions are asynchronous, a second division of two daughters occurs within a maximum interval of 20 minutes. During the mid-gastrulation and early segmentation periods, we established that the cell cycle length of r-progenitors is 4hpf35 +/-20 minutes maximum on average (Fig. 3C; 4h55 +/-25 minutes in a second data set ID190828aZ, Fig. S1C). Similarly, a previous study reported that the cell cycle length of CNS progenitors at division 15 (8-10hpf) and division 16 (10-15hpf) averages 151+/-59 and 240+/-71 minutes, respectively (Kimmel, Warga and Kane, 1994). There is no difference in the proliferation rate for individual rhombomeres or in dorsal versus ventral progenitor populations (Fig. 3C and D, Fig. S1C). We analyzed the spatial distribution of dividing cells and found this to be uniform throughout the neuroectoderm, from shield to early neurula stages (Fig. 3E). This uniform spatial pattern of cell division was also reported during xenopus neurulation (Christodoulou and Skourides, 2022). The reconstruction of the cell lineage trees of rhombomeres’ r2-r6 establish that the clonal commitment of rhombomeres is set by mid-gastrulation and that the segments are mainly formed by proliferation with a uniform proliferation rate along the AP axis of hindbrain and little intermingling of cells between adjacent segments.

**Fig. 3.**
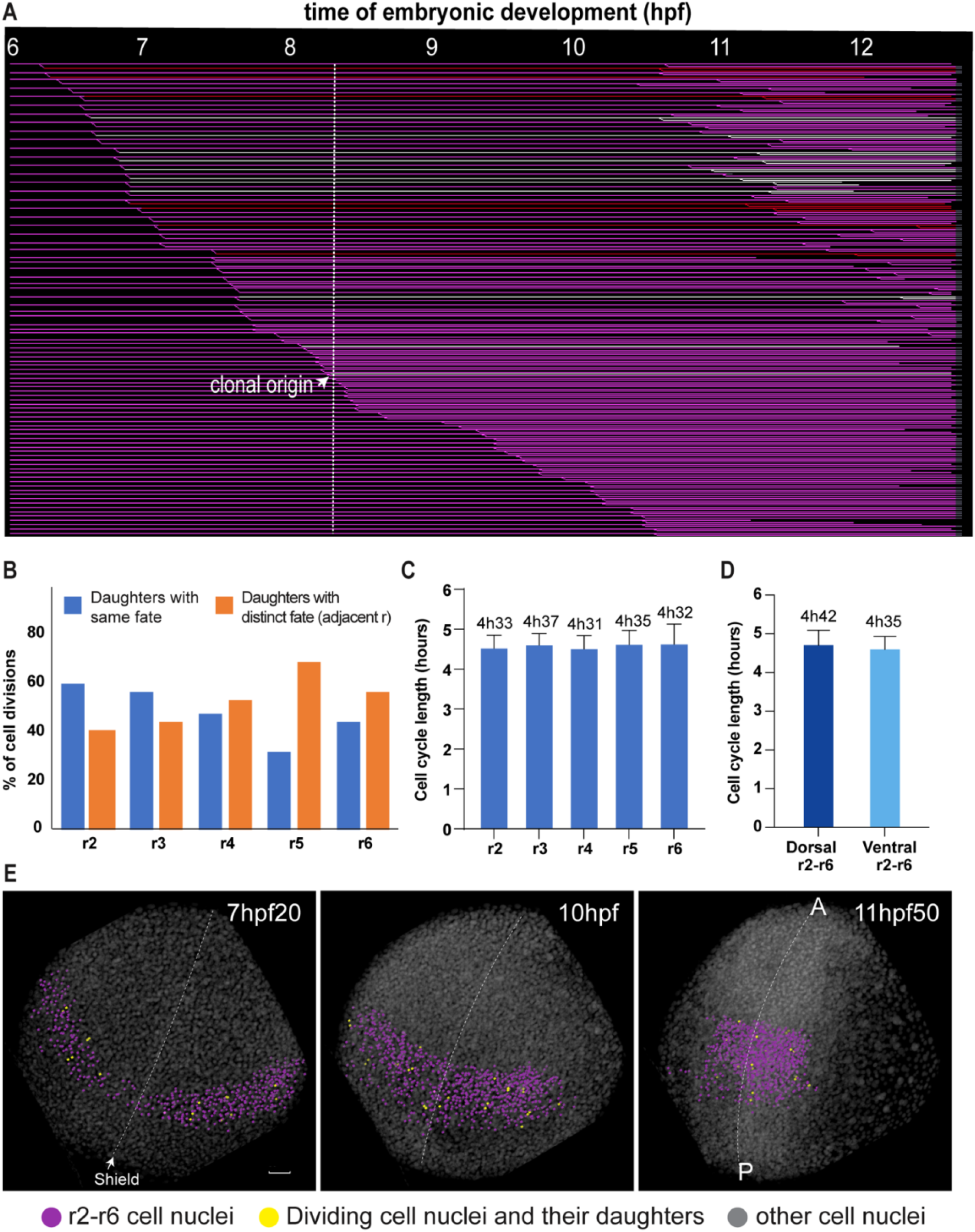
Pattern of cell division sets the clonal origin of r3 by mid gastrulation. A) Flat representation of the reconstructed cell lineage tree of r3 progenitors from 6hpf to 12hpf40 (x axis) displayed according to the timing of the first observed division along the lineage. Each line represents a single hindbrain progenitor. Some cells were not tracked until the end of the sequence leading to interrupted lines. The clonal origin of r3 is set at 8hpf20 (white dotted line). Arrow indicates the last cell division giving rise to daughters fated to different (adjacent) rhombomeres. r3 pink, r2 red, r4 white. B) Percentage of cell divisions leading to daughters with identical or distinct (adjacent rhombomeres) fates before the clonal origin. C) Cell cycle length for r2-r6 progenitors between 6 to 13hpf. Number of divisions analyzed: r2 61, r3 55, r4 60, r5 45, r6 25. D) Cell cycle length of r2-r6 dorsal and ventral progenitors from 6 to 13hpf. Number of divisions analyzed: dorsal 42, ventral 66. E) Spatial distribution of cell divisions, mother or daughter cells (yellow) in the hindbrain (nuclei in violet), nuclei raw data in gray, snapshots at 7hpf20, 10hpf,11hpf50, 3D rendering, dorsal view, dotted white line for the midline; A: anterior, P: posterior. White arrow indicates the shield position. Scale bar 50 µm.

### Summary and discussion

We developed a method to study zebrafish hindbrain morphogenesis quantitatively from mid-gastrulation till early neurulation based on 3D+time live imaging of a transgenic embryo with a hindbrain-specific reporter. Technical advances in two-photon scanning microscopy and automated cell tracking tools enable such a cellular approach. Quantifying cell behavior in terms of proliferation and movement will serve as a basis to build a mechano-genetic model of hindbrain formation *in silico*. Hindbrain rhombomeres start to be specified as early as 8hpf as evidenced by marker gene expression. Thus, this approach can be used to understand the early cell behaviors that lead to hindbrain defects upon gain and loss of various gene functions in the course of hindbrain morphogenesis.

We provided a zebrafish hindbrain fate map with improved precision compared to the state-of-the-art (Woo and Fraser, 1995) by exploiting technical advances (Fig. 8). Our fate map shows no dramatic wavy domains of hindbrain progenitors (highlighted in violet) as predicted in (Woo and Fraser, 1995). We tracked several hundred cells including those in deep layers in the hindbrain compared to the smaller number of surface-accessible cells analyzed in earlier work. However, the r1 and r7 domains were not included in the analysis of our fate map. This may result in regions that overlap with midbrain and spinal cord progenitors at the anterior and posterior ends respectively. Single-cell resolution of neural progenitors (neuroectoderm) shows that they undergo orderly movement throughout gastrulation. Progenitor populations of each rhombomere are organized segmentally as early as the shield stage (Fig. 8A). A closer examination showed about 12-18% overlap with progenitor populations of flanking rhombomeres’. This overlap resolves over time and thus forms sharp boundaries at later stages. Dorsal and ventral progenitors of the hindbrain are located more laterally and medially, respectively, relative to the embryonic shield. Both populations move in the same direction towards the midline. At 10hpf30, these medially located cells converge on the midline and follow anterior and inward movement to achieve internalization (Fig. 8B). Lateral cells located far from the midline accelerate and converge directly to the midline. They thus place themselves on top of inward-moving medial cells. Dorsal and ventral progenitors undertake distinct migration paths to orchestrate the 3D neural keel structure starting from the 2D neural plate. Previous study covered the interval from late gastrulation to early neurulation and reported the differential migration paths of dorsal and ventral progenitors at the posterior hindbrain/spinal cord levels (Araya *et al*., 2019). We were able to capture and examine cell behaviors that underlie the transition from neuroectoderm to neural keel with continuous coverage at single-cell resolution. The molecular and cellular mechanisms underlying the differential migration paths of dorsal and ventral progenitors during early neurulation must be determined. The observed increase in velocity of lateral cells in our study was reported previously using conventional methods but did not provide the exact time period (Sepich *et al*., 2000; Myers, Sepich and Solnica-Krezel, 2002). We demonstrated that clonal commitment of rhombomere progenitors is achieved at around 8hpf-9hpf. The defined clonal origin of a rhombomere is in agreement with transplantation experiments showing that regional commitment to neural identity occurs at 80% epiboly (Woo and Fraser, 1998). Divisions of progenitors give rise to daughters of only same or adjacent rhombomeres but not of other rhombomeres. It indicates that the cell mixing between rhombomeres is limited. The cell cycle length of neural progenitors seems to vary slightly between two embryos, 4h30min +/-20min in one and 4h55 +/-25 minutes in another. We further showed that the spatial and temporal patterns of cell proliferation are uniform during mid-gastrulation through early neurulation. Overall, our study provided precision to previous work performed through classical methods and further dissected the cell behaviors of hindbrain progenitors quantitatively and provided a dynamic fate map of the zebrafish hindbrain at single-cell resolution.

**Fig. 8.**
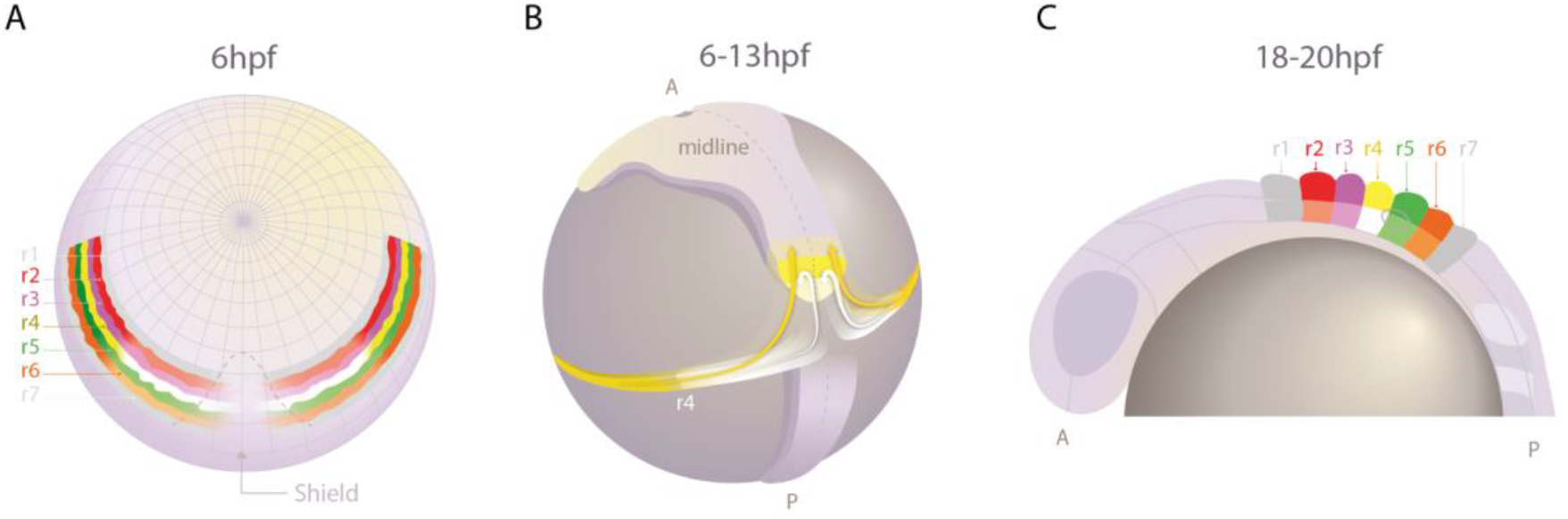
Dynamics of the zebrafish hindbrain fate map throughout gastrulation and neurulation. A) Rhombomere’ progenitors from r2 to r6 are labeled with their respective color code, r1 and r7 are labeled arbitrarily at the shield stage. Slight wavy borders show the cell intermingling at the rhombomere borders. Ventral progenitors at the shield stage are marked in light colors and their respective dorsal counterparts in dark colors except that r4 dorsal in yellow and r4 ventral in white. B) Mean trajectory of the dorsal (yellow) and ventral (white) progenitors of r4 from 6 to 13hpf on 13hpf embryo. Dorsal and ventral progenitors take distinct paths to form the neural tube. C) Neural rod with r2-r6 segments with their respective color code. r1 and r7 are labeled in light gray. Dorsal parts are labeled with dark colors and their ventral counterparts are labeled with light colors. D dorsal, V ventral, A anterior, P posterior.

## Material and Methods

### Zebrafish transgenesis and husbandry

Transgenic fish lines, *krox20:eGFP-Hras* were generated to label developing rhombomere domains r3 and r5 with eGFP targeted to membranes. A SalI/ClaI fragment containing GFP-Hras from pCS2/GFP-Hras was subcloned into pCR2.1-TOPO to be transferred as a BamHI/NotI fragment in pTol2-cA:GFP construct (Labalette *et al*., 2011). Transgenic lines were obtained from WT/Tü and *casper* embryos injected at the one-cell stage with the pTol2 constructs together with tol2 transposase mRNA. The zebrafish lines were bred and maintained as described (Westerfield, 2007). Embryos were staged in accordance with (Kimmel *et al*., 1995).

### mRNA preparation and microinjection

*H2B-mCherry* and *nls-Eos* mRNAs were prepared using mMessage mMachine mRNA transcription kit (Invitrogen) and stored in -80°C. 3-4 nl of 75 ng/µl H2B-mCherry mRNA was injected in one-cell stage zebrafish embryos. Injection at an early one-cell stage into the center of the cell of the embryo ensured homogenous staining. Embryos with homogenous and optimum bright staining were selected either around 4hpf for live imaging experiments.

### Mounting for embryo at 6hpf for imaging hindbrain formation

Embryo at 6hpf was dechorionated and mounted in the custom-made imaging chamber on a bed of 0.5% LMP Agarose. The imaging chamber was filled with EM. Tricaine was used at 0.033% in EM to inhibit twitching of embryos at later stages. First, the embryo was placed on an agarose bed with animal pole on top and then the shield position is tilted up to 30º angle (1/3^rd^ to the center of the embryo) and immobilized by pouring 20-30 μl of 0.5% LMP agarose on top. This initial position of the shield stage embryo ensured that the hindbrain progenitors were inside the field of view throughout the gastrulation and neurulation movements.

### Two-photon and confocal microscopy imaging

Leica SP5 and Zeiss LSM780 two-photon microscopes equipped with water immersion 20X/1.0 NA dipping lens objectives were used for overnight imaging. Image stacks of 200-250 μm with spatial scaling 1-1.16 μm and with delta T ∼2m30s were made continuously for 10 hours and more. While starting the imaging, 70μm was scanned above the animal pole to ensure the embryo’s growth inside the volume scanned during the imaging period. Embryos were maintained at 28.5°C with the use of an OKO-lab heating system (OKO-lab H101 WJC). Only embryos which exhibited normal morphology after imaging were analyzed. Dechorionated embryos were mounted in customized agarose gel molds. Image stacks 100-120 μm were made at 1024×1024 spatial resolution.

| ID dataset | Transgenic & Background | Time per stack | Imaging period |
| --- | --- | --- | --- |
| 180516aF | Krox20:eGFP-ras, Casper | 2m27s | 6hpf-20hpf |
| 190828aZ | Krox20:eGFP-ras, Casper | 2m33s | 6hpf-20hpf |

### Data Analysis

Raw data was processed through BioEmergences workflow to obtain the automated reconstruction of cell lineages (Faure *et al*., 2016). Nuclei centers were validated through center-select software. A nuclei-based tracking algorithm called Expectation-Maximization (EM), which is the iteration of the SimAnn algorithm (published previously from the lab) was applied. The raw data and reconstructed digital embryos were visualized, annotated and analyzed in Mov-IT, a custom made interface (Faure *et al*., 2016). Parameters such as mean trajectory, cell cycle length and velocity were extracted from Mov-IT software. Graphpad prism and MS excel were used for statistical analysis and representation. Mann-whitney and student t-tests were performed to find the significance of compared groups. Images for the publication were made using the same software. Fiji software was used to analyze and present confocal images. Figures were assembled in Adobe illustrator.

## Acknowledgements

We thank Mark E. Hammons and Maxime Comberiati for the assistance with Bio-Emergences workflow and fish facility respectively. We thank Dr David Bensimon and Dr Paula Alexandre for their valuable comments. We acknowledge Olga Markova for the graphical illustration. We thank ImageInLife, European Union’s Horizon 2020 research and innovation programme under the Marie Sklodowska-Curie grant agreement No. 721537 for providing three year PhD funding to MK (2017-2020) and Graduate School Life sciences and Health, University Paris-Saclay for providing three months funding to MK.

---

**Movie 1. Dynamic fate map of hindbrain (r2-r6) from mid-gastrulation till early neurulation**. Rhombomere progenitors’ migration from 6hpf till 14hpf20. Color code: r2 red, r3 pink, r4 white, r5 green, r6 orange, other raw nuclei in gray. r2-r4 progenitors’ nuclei were not tracked beyond 12hpf40. Dorsa-lateral view. Scale bar 50 µm.

http://bioemergences.eu/MageshiK/videos/180516aF_annotated.mp4

**Movie 2. Construction of dorsal and ventral 3D neural keel (r4) from progenitors domains**. r4 dorsal and ventral progenitors’ migration from 6hpf till ∼13hpf. Color code: r4 dorsal in yellow and r4 ventral in white, labeled on the right side of the embryo, other raw nuclei in gray. Both r4 dorsal and ventral progenitors are labeled white on the left side. Dorsal view. Scale bar 50 µm.

http://bioemergences.eu/MageshiK/videos/180516aF_t_t21383_26.mp4

**Movie 3. Construction of 3D neural keel from neuroectoderm through transverse view**. r4 dorsal and ventral progenitors’ migration from 7hpf till ∼14hpf. Color code: r4 dorsal in yellow and r4 ventral in white labeled on the right side, other raw nuclei in blue. Transverse view. Scale bar 20 µm.

http://bioemergences.eu/MageshiK/videos/180516aFDVformation.mp4

