## Supplementary figures for "Dynamic fate map of hindbrain rhombomeres in zebrafish"

**Fig. S1.**

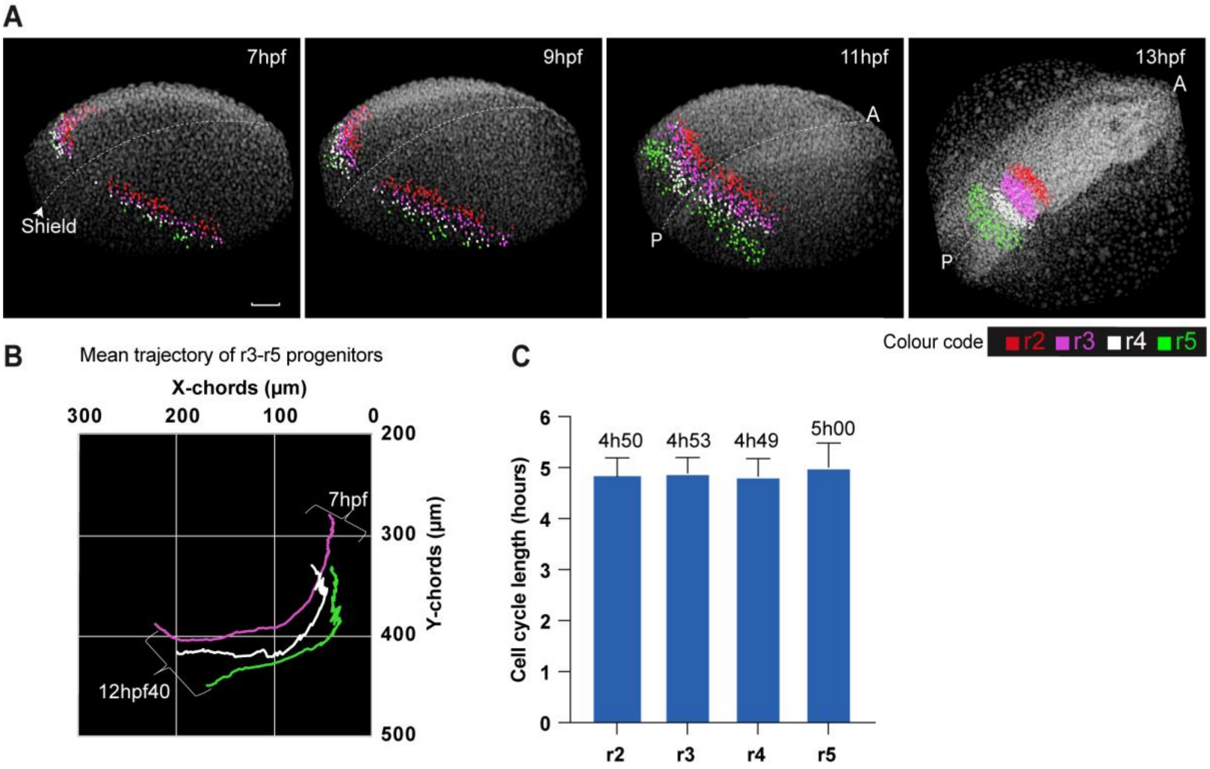


**Segmental organization of rhombomeres' progenitors is refined during gastrulation and early neurulation.** Second dataset ID: 190828aZ A) Analysis of the processed data, dorso-lateral view 3D rendering with nuclei staining raw data in gray, anterior top right (A), posterior bottom left (P), scale bar 50 µm, midline marked by a dotted white line. Color code of the rhombomeres is given at the bottom. r2-r5 progenitors' nuclei position at 7hpf, 9hpf, 11hpf, 13hpf. B) Mean trajectory of the r2, r3, r4 and r5 progenitor population (one side of the embryo) is plotted in the XY axis from 7hpf till 13hpf. Note that the number of cells manual corrected are less compared to the first dataset. Therefore, the trajectories of the rhombomeres are not perfectly parallel C) Cell cycle length of r2, r3, r4 and r5 progenitors during the embryonic period, 6 to 13hpf. Number of divisions analyzed: r 42, r3 48, r4 29, r5 9.

**Fig. S2.**

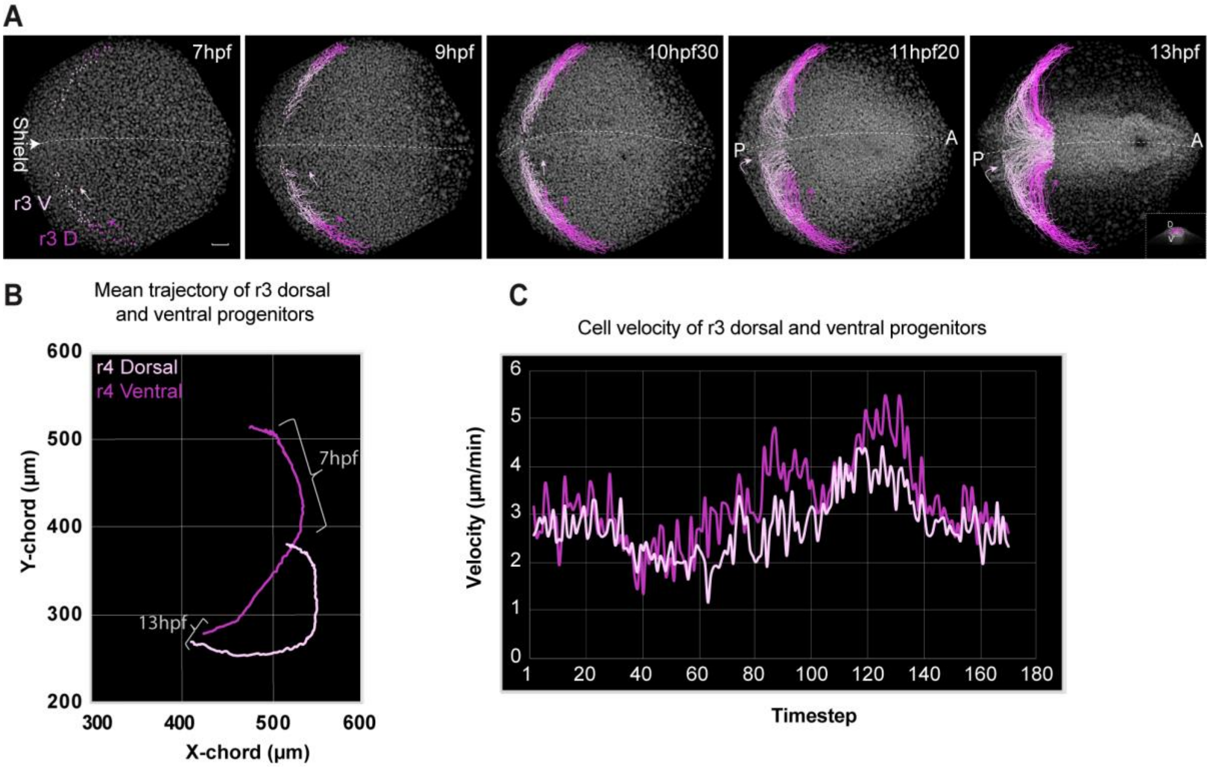


**Dorsal and ventral hindbrain progenitors have distinct convergence/extension paths.** Second dataset ID: 190828aZ A) 3D rendering of image data, nuclei raw data in gray, nuclei selected with color codes for dorso-ventral domains, time in hpf top left, midline indicated in white dotted, orientation anterior to the left, scale bar 50μm. r4 progenitors' position and trajectory from 7hpf to 13hpf, dorsal (D pink) and ventral (V white), arrowhead points to the shield, pink and white arrows indicate the direction of progenitor migration. A anterior P posterior. D dorsal. V ventral. B) Mean trajectory of dorsal and ventral progenitors of rhombomere 3 (one side of the embryo) is plotted in the XY axis from 7hpf till 13hpf. C) Relative velocity of dorsal and ventral progenitors of r3 from 6hpf to 13hpf (Timestep 1-180).

**Fig. S3.**

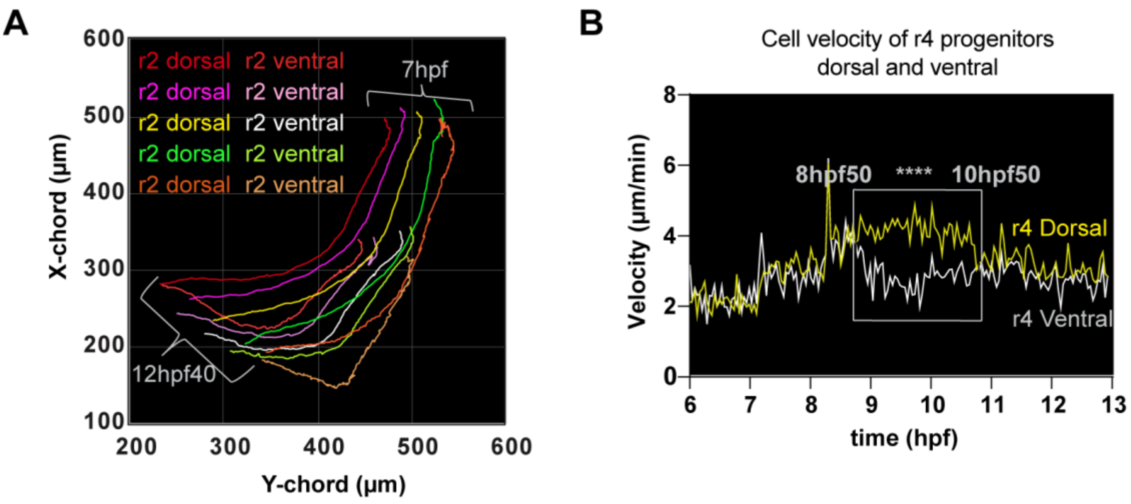


**DV trajectories and cell velocity of rhombomeres.** First dataset ID: 180516aF A) Mean trajectory of dorsal and ventral progenitors of each rhombomere population r2 in dark and light red, r3 in dark and light pink, r4 in yellow and white, r5 in green and light green, r6 in orange and light orange respectively from 7hpf till 12hpf40 is plotted in the XY axis. B) Relative velocity of dorsal (yellow) and ventral (white) progenitors of r4 from 6hpf till 13hpf. P=5.8171823800557726e-12.

**Fig. S4.**

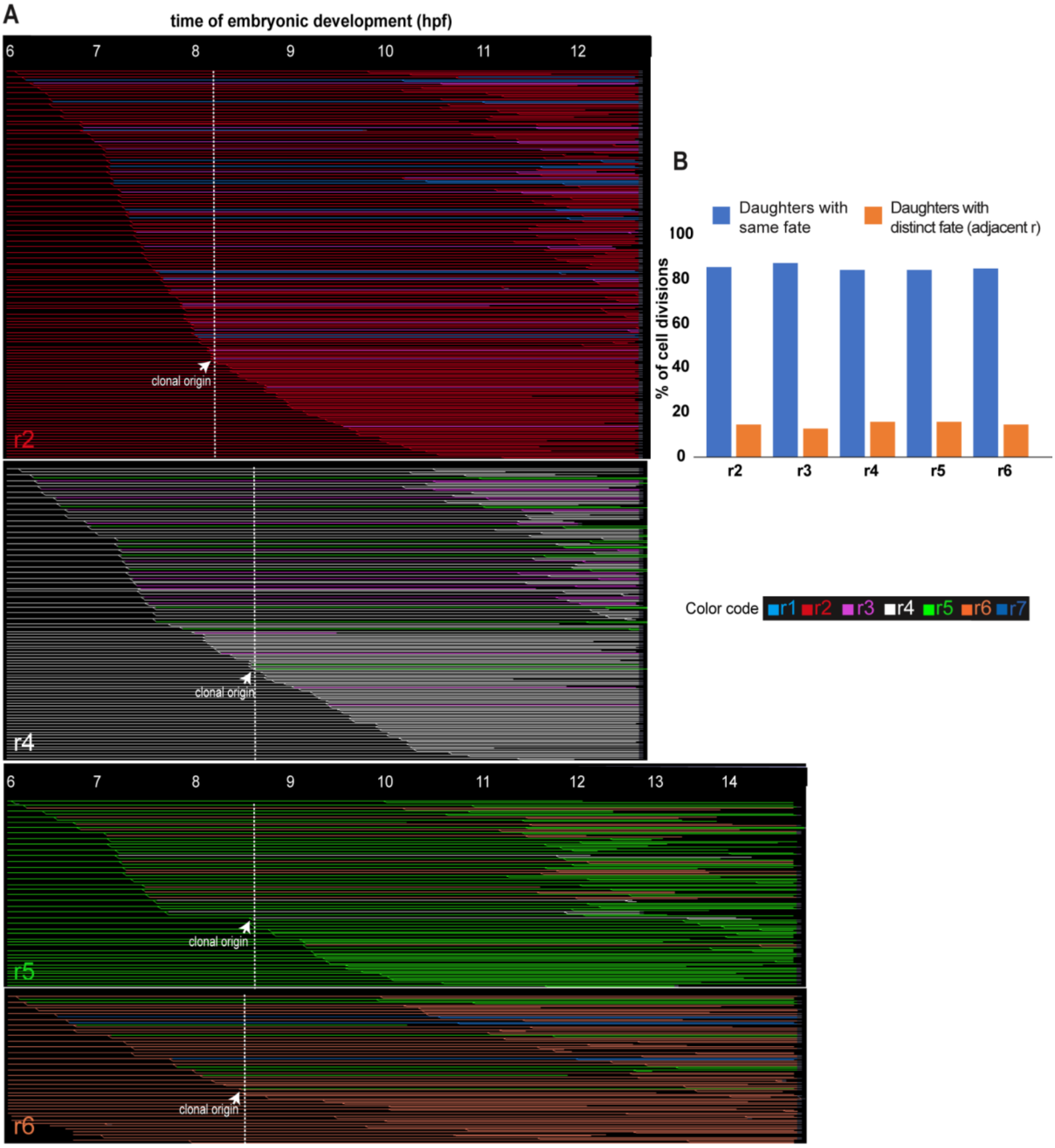


**Cell lineage trees of rhombomeres r2- r6.** First dataset ID: 180516aF A) Flat representation of the reconstructed cell lineage tree of r3 progenitors from 6hpf to 12hpf40 (x axis) displayed according to the timing of the first observed division along the lineage. Each line represents a single hindbrain progenitor. Some cells were not tracked until the end of the sequence leading to interrupted lines. The clonal origin in each lineage tree is indicated with a white dashed line. Arrow indicates the last cell division giving rise to daughters of different (adjacent) rhombomere fate. r2 at 8hpf14, r4 at 8hpf38, r5 at 8hpf36, r6 at 8hpf29. Color code is given at the right. r2 red, r3 pink, r4 white, r5 green and r6 orange. Blue lines in r2 and r6 lineage trees indicate r1 and r7 respectively. B) Percentage of cell divisions leading to daughters with identical or distinct (adjacent rhombomeres) fates analyzed from 6hpf to 14hpf.
